# Who bears the load? IOP-induced collagen fiber recruitment over the corneoscleral shell

**DOI:** 10.1101/2022.12.10.519931

**Authors:** Tian Yong Foong, Yi Hua, Rouzbeh Amini, Ian A. Sigal

## Abstract

Collagen is the main load-bearing component of cornea and sclera. When stretched, both of these tissues exhibit a behavior known as collagen fiber recruitment. In recruitment, as the tissues stretch the constitutive collagen fibers lose their natural waviness, progressively straightening. Recruited, straight, fibers bear substantially more mechanical load than non-recruited, wavy, fibers. As such, the process of recruitment underlies the well-established nonlinear macroscopic behavior of the corneoscleral shell. Recruitment has an interesting implication: when recruitment is incomplete, only a fraction of the collagen fibers is actually contributing to bear the loads, with the rest remaining “in reserve”. In other words, at a given intraocular pressure (IOP), it is possible that not all the collagen fibers of the cornea and sclera are actually contributing to bear the loads.

To the best of our knowledge, the fraction of corneoscleral shell fibers recruited and contributing to bear the load of IOP has not been reported. Our goal was to obtain regionally-resolved estimates of the fraction of corneoscleral collagen fibers recruited and in reserve. We developed a fiber-based microstructural constitutive model that could account for collagen fiber undulations or crimp via their tortuosity. We used experimentally-measured collagen fiber crimp tortuosity distributions in human eyes to derive region-specific nonlinear hyperelastic mechanical properties. We then built a three-dimensional axisymmetric model of the globe, assigning region-specific mechanical properties and regional anisotropy. The model was used to simulate the IOP-induced shell deformation. The model-predicted tissue stretch was then used to quantify collagen recruitment within each shell region. The calculations showed that, at low IOPs, collagen fibers in the posterior equator were recruited the fastest, such that at a physiologic IOP of 15 mmHg, over 90% of fibers were recruited, compared with only a third in the cornea and the peripapillary sclera. The differences in recruitment between regions, in turn, mean that at a physiologic IOP the posterior equator had a fiber reserve of only 10%, whereas the cornea and peripapillary sclera had two thirds. At an elevated IOP of 50 mmHg, collagen fibers in the limbus and the anterior/posterior equator were almost fully recruited, compared with 90% in the cornea and the posterior sclera, and 70% in the peripapillary sclera and the equator. That even at such an elevated IOP not all the fibers were recruited suggests that there are likely other conditions that challenge the corneoscleral tissues even more than IOP. The fraction of fibers recruited may have other potential implications. For example, fibers that are not bearing loads may be more susceptible to enzymatic digestion or remodeling. Similarly, it may be possible to control tissue stiffness through the fraction of recruited fibers without the need to add or remove collagen.

## 1. Introduction

Collagen fibers are the main load-bearing component of the eye. (Boote et al., 2020) From simple fiber embedded models (Amini and Barocas, 2009) to more complex ones (Grytz and Meschke, 2010), the anisotropy and microstructural architecture of ocular collagen fibers have been shown to significantly affect the biomechanics of the corneoscleral shell. Hence, great efforts have been devoted to developing imaging tools for mapping collagen fibers. (Campbell et al., 2015; Danford et al., 2013; Girard et al., 2011; Ho et al., 2014; Jan et al., 2015; Ling et al., 2019; Pijanka et al., 2019; Winkler et al., 2011; Zhang et al., 2015; Zhou et al., 2019b) The fiber maps are then translated into mechanical models. The underlying assumption being that the collagen maps directly translate into the local capability of the tissues to bear load. (Grytz and Meschke, 2010) However, as we, (Gogola et al., 2018a; Jan et al., 2018; Jan et al., 2017a) and others, (Boote et al., 2020; Liu et al., 2014) have shown, the tissues of the eye exhibit natural undulations called crimp. Wavy fibers, in turn, lead to an important behavior of the tissues: recruitment. When under load, the initially wavy fibers straighten. While the wavy fibers bear almost no load, the straightened ones bear substantial loads. The process of progressively straightening fibers is known as fiber recruitment (**Figure 1**). Fiber recruitment has been shown to underlie the nonlinear mechanical behavior of many soft tissues, not only sclera and cornea. (Cheng et al., 2018; Fata et al., 2014; Hansen et al., 2002; Hill et al., 2012; Jan and Sigal, 2018; Mattson et al., 2017; Roy et al., 2010; Weisbecker et al., 2015)

**Figure 1.**
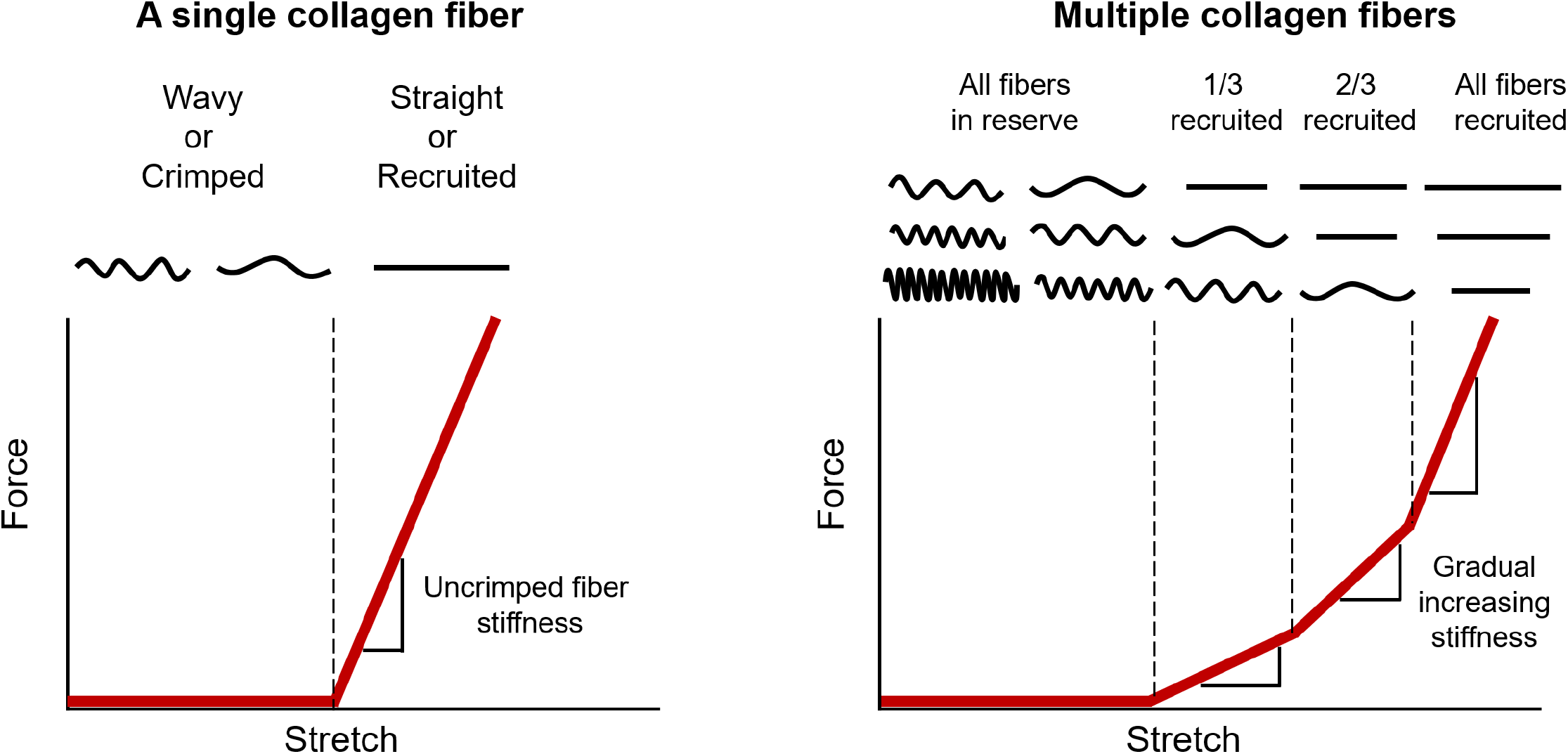
Diagram of collagen fiber recruitment. **(Left)** A single fiber is easy to stretch when it is crimped or wavy. However, once the fiber is straightened or recruited, the fiber becomes much stiffer, requiring more force to elongate the fiber. **(Right)** When many crimped fibers with different amount of slack are stretched together, the gradual straightening of the fibers creates a gradual, nonlinear stiffening, where more and more force is required to continue stretching the tissue. A fiber that is straightened is referred to as recruited. Conversely, a fiber that is not recruited, but would at higher stretch, is referred to as in reserve. The proportion of fibers recruited to those in reserve is directly related to the rate of recruitment. Adapted from (Jan and Sigal, 2018).

During the process of recruitment, only a fraction of tissue collagen fibers are load-bearing, while the rest remains in “reserve”. In other words, similar to the dormancy of cells, the undulated fibers are “off”. When straightened, the fibers turn “on” and are considered recruited to bear mechanical loads. For example, we have recently shown that at normal IOP, only about three-quarters of the fibers of the peripapillary sclera and lamina cribrosa in sheep eyes are contributing to bearing the load, with one-fourth in reserve. (Jan and Sigal, 2018) Peripapillary sclera and lamina cribrosa had distinct recruitment curves, with different fractions of fibers recruited at sub-physiologic and supra-physiologic pressures. With the rest of the corneoscleral shell exhibiting the same fundamental mechanism of recruitment, it follows that at normal pressures also only a fraction of the fibers is contributing to bear the load of IOP. This can have important implications. First, from a fundamental perspective it is interesting and worthwhile to note that it is not necessarily correct to assume that collagen fibers will help bear a load simply because they “are there”. Hence, fibers discernible in stained sections or by other imaging tools, may or may not contribute to bear a particular load. Second, the recruitment may also have important implications for remodeling and treatments aimed at modifying tissue properties. For instance, fibers not bearing load are likely to be more susceptible to degradation, whereas fibers under load are more likely to remodel. Similarly, fiber cross-linking likely affects fibers differently depending on whether they are bearing load or not. Moreover, given the regional variations in micro and macrostructure and mechanics over the globe, (Gogola et al., 2018a; Jan et al., 2018; Voorhees et al., 2017a; Whitford et al., 2016) it seems reasonable to expect that the fraction of recruited fibers likely also varies. Further, the fraction of fibers in reserve when under load can indicate the extent to which a region is able to adapt to further loads or if it has reached a limit.

Despite the abovementioned reasons, to the best of our knowledge, there is no information on the fraction of corneoscleral fibers bearing load at a given IOP. Our goal in this study was to obtain regionally-resolved estimates of the fraction of corneoscleral collagen fibers recruited, hence “turned-on” for bearing IOP-related loads. To achieve this goal, we used computational modeling based on experimentally obtained measures of collagen fiber undulations. (Gogola et al., 2018a) As a first step, in this work, we focused on estimating the fiber recruitment fraction in young healthy eyes.

## 2. Methods

Our general strategy was as it follows (**Figure 2**): using polarized light microscopy (PLM), we measured collagen crimp tortuosity in seven regions across the corneoscleral shell. The results were then fed into a fiber-based microstructural constitutive model to obtain region-specific nonlinear hyperelastic mechanical properties. A three-dimensional axisymmetric model was developed to simulate the IOP-induced deformation of the corneoscleral shell. The model-predicted tissue stretch was used to quantify collagen fiber recruitment over the corneoscleral shell. Below we describe each step in detail.

**Figure 2.**
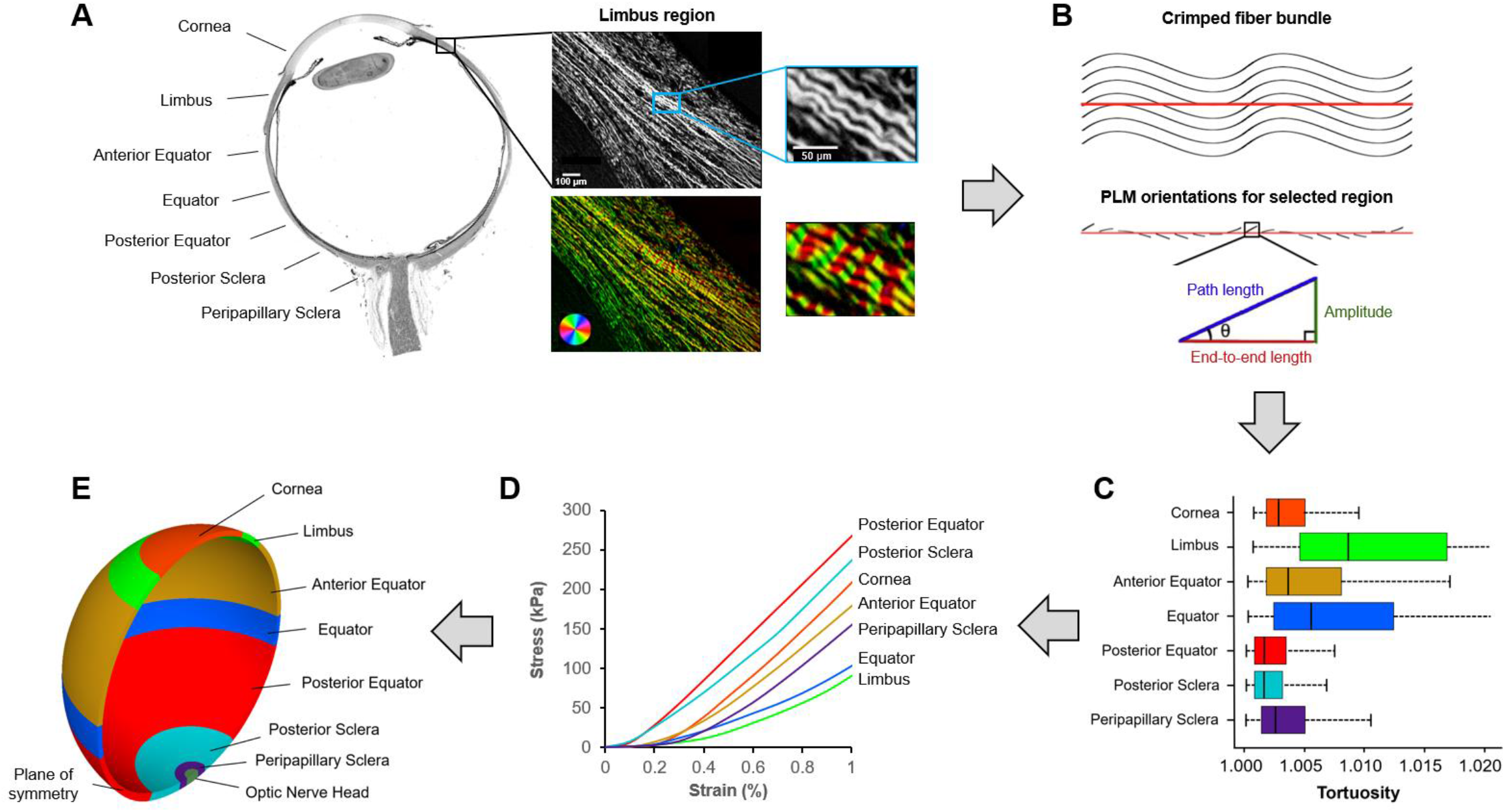
General strategy. We divided the corneoscleral shell into seven regions **(A)**. Each region was imaged using polarized light microscopy (PLM). The close-up images show the collagen fibers in the limbus region. The wavy pattern of collagen fibers is discernible and emphasized using a color map of orientation. To quantify collagen crimp, a straight line was marked manually along a fiber bundle **(B)**. Trigonometric identities were then used pixel by pixel along the line to calculate local amplitude, path length, and end-to-end length based on the orientation information derived from PLM as described in detail in (Brazile et al., 2018; Gogola et al., 2018a; Jan et al., 2018). The local amplitude and lengths were then integrated to compute those of the fiber bundle. The crimp tortuosity of the fiber bundle was calculated as its integrated path length divided by the end-to-end length. We measured collagen crimp tortuosity in seven regions across the corneoscleral shell and the results **(C)** were fed into a fiber-based microstructural constitutive model to obtain region-specific nonlinear hyperelastic mechanical properties **(D)**. A three-dimensional axisymmetric model **(E)** was then developed to simulate the IOP-induced deformation of the corneoscleral shell. The model-predicted tissue stretch was used to quantify collagen fiber recruitment over the corneoscleral shell.

### 2.1 Measuring collagen crimp across the corneoscleral shell: a review

Details of sample preparation, imaging, and crimp quantification were given in our previous report. (Gogola et al., 2018a) A brief description of these methods are provided here:

#### Sample preparation

Nine normal eyes of nine human donors (ages ranging from 1 month to 17 years) were fixed unpressurized (0 mmHg IOP) in 10% neutral buffered formalin for a week. The eyes were then bisected into superior and inferior portions without cutting through the optic nerve head. The superior portion, which contained the optic nerve head, was embedded in paraffin and cryosectioned axially at 5-µm thickness. A total of 42 sections, which passed through both the optic nerve head and the cornea and were free of artifacts, such as tears and folds, were selected for imaging.

#### Imaging

The selected sections were imaged with PLM using a previously reported method to visualize and quantify collagen fiber orientations. (Brazile et al., 2018; Gogola et al., 2018b; Jan et al., 2017a; Jan et al., 2015; Jan et al., 2017b) For each section, images were collected from seven regions across the corneoscleral shell: the cornea, limbus, anterior equator, equator, posterior equator, posterior sclera, and peripapillary sclera (**Figure 2A**).

#### Crimp quantification

Crimp tortuosity was measured in at least 20 collagen fiber bundles in each region, distributed throughout depth (**Figure 2B**). For each bundle, we manually placed a line segment with a length of 60 pixels (∼ 45 µm). This line segment was used to sample the collagen fiber orientation values within the bundle for calculating the crimp tortuosity. We refer readers to our previous papers for full details of the calculation of crimp tortuosity. (Brazile et al., 2018; Gogola et al., 2018a; Jan et al., 2018) The distribution of crimp tortuosity in the seven regions across the corneoscleral shell is shown in **Figure 2C**.

### 2.2 Crimp-informed nonlinear hyperelastic mechanical properties

For tissue consisting of fibers with various tortuosities, the gradual straightening of the fibers creates a nonlinear stiffening. (Brazile et al., 2018; Cacho et al., 2007; Jan and Sigal, 2018) To obtain the region-specific mechanical behavior of the corneoscleral shell, we pooled crimp tortuosity measurements in each region and then calculated the corresponding stress-strain curve (**Figure 2D**). Specifically, the mechanical response of the corneoscleral shell was modeled using a fiber-based constitutive equation of the form

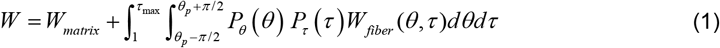

where *W* is the total strain energy density; *W*_*matrix*_ is the isotropic strain energy density of the ground substance; *W*_*fiber*_ (*θ, τ*) is the strain energy density of the anisotropic fibers; *P*_*θ*_(*θ*) and *P*_*τ*_(*τ*) are the probability density functions used to describe the percentage of fibers with a given orientation and tortuosity, respectively; *θ*_*p*_ is the preferred fiber orientation relative to a local coordinate system. *τ* is the tortuosity of a crimped fiber (≥ 1); *τ*_*max*_ is the maximum tortuosity in a specific region of the tissue.

The strain energy density equation for the ground substance was modeled as a Neo-Hookean material which has the form

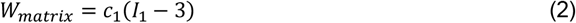

where *I*_1_ is the first invariant of the right Cauchy-Green deformation tensor; *c*_1_ is the first Mooney-Rivlin coefficient, which is also equivalent to the matrix shear modulus divided by two. We chose *c*_1_ to be 150 kPa. (Girard et al., 2009a; Hua et al., 2020; Voorhees et al., 2017c)

As shown in Figure 1, we assumed a fiber is easy to stretch when it is crimped or wavy, and its stress is zero; once the fiber is straightened, it starts to bear load, and its stress increases linearly with stretch. (Brazile et al., 2018; Cacho et al., 2007; Jan and Sigal, 2018) Based on this assumption, the strain energy associated with a single collagen fiber at a given orientation and tortuosity satisfies the following relationship

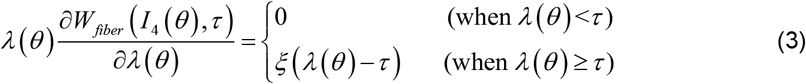

where *I*_4_(*θ*) is the fourth invariant of the right Cauchy-Green deformation tensor associated with the collagen fiber family aligned in the orientation *θ*, which is equivalent to the squared fiber stretch *λ*(*θ*); *τ* is the tortuosity of a crimped fiber (≥ 1); *ξ* is the elastic modulus of a straightened fiber, which was set to 30 MPa. (Grytz and Meschke, 2009)

We then obtained the Cauchy stress tensor as follows

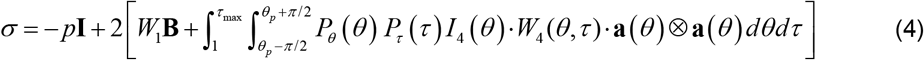

where *p* is the Lagrange multiplier to enforce incompressibility (similar to the hydrostatic pressure in fluids); **I** is the second-order identity tensor; **B** is the left Cauchy-Green deformation tensor; **a** is a unit vector representing the local fiber direction in the deformed configuration. *W*_1_ and *W*_4_ are expressed as

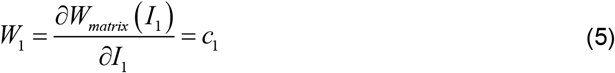

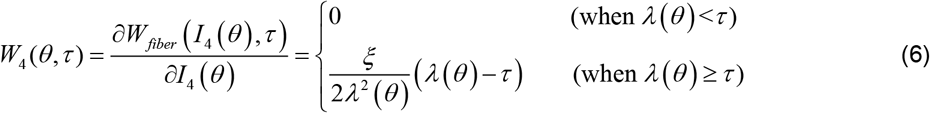

We checked the convexity requirements for the proposed constitutive model to ensure its reliable mechanical and mathematical behavior (Supplementary Material).

### 2.3 Finite element modeling

#### Model geometry

To reduce computational cost, we modeled the only one hemisphere of the corneoscleral shell with a simplified optic nerve head (**Figure 2E**). The inner radius of the shell was 12 mm. (Sigal et al., 2005a; Sigal et al., 2004) The shell thickness was non-uniform, with a maximum thickness of 996 µm adjacent to the scleral canal and a minimum thickness of 491 µm at the equator. (Norman et al., 2010; Norman et al., 2011) A spline fit was performed to ensure a smooth transition in thickness. (Wold, 1974) The shell was then partitioned circumferentially into seven regions, matching those in the experiment. (Gogola et al., 2018a)

#### Mechanical properties

An anisotropic hyperelastic constitutive equation was used to describe the region-specific mechanical behavior of the corneoscleral shell, (Girard et al., 2009b) with parameters determined by fitting the crimp-informed nonlinear stress-strain curves (**Figure 2D**). The collagen fibers of the corneoscleral shell followed the semi-circular von Mises distribution. In the limbus and the peripapillary sclera, the collagen fibers were aligned circumferentially; while in the cornea, we defined two preferred collagen fiber orientations orthogonal to each other. (Abahussin et al., 2009) The collagen fibers in the other regions of the corneoscleral shell were assumed to be randomly distributed. A bulk modulus of 0.1 GPa was defined to ensure tissue incompressibility. (Girard et al., 2009b) Note that the bulk modulus does not directly correspond to the tissue compressibility. Instead, it acts as a penalty factor that helps define how strict the incompressibility constraint is enforced. The value selected provides reliable model convergence while keeping volume changes low. A summary of the hyperelastic parameters is listed in **Table 1**.

**Table 1.** Region-specific hyperelastic parameters used in finite element modeling. Please see the main text for more details of how the parameters were obtained.

| | Mooney-Rivlin parameter<br>$c_1$ (kPa) | Exponential multiplier<br>$c_3$ (kPa) | Fiber scale factor<br>$c_4$ (-) | Fiber concentration factor<br>$k_f$ (-) |
| --- | --- | --- | --- | --- |
| Cornea | 150 | 39 | 184 | 2 |
| Limbus | 150 | 6 | 275 | 2 |
| Anterior equator | 150 | 35 | 181 | 0 |
| Equator | 150 | 16 | 196 | 0 |
| Posterior equator | 150 | 232 | 76 | 0 |
| Posterior sclera | 150 | 135 | 100 | 0 |
| Peripapillary sclera | 150 | 15 | 240 | 2 |

The optic nerve head was modeled as an incompressible, linear isotropic material with an elastic modulus fixed to 0.3 MPa. (Sigal et al., 2005a; Sigal et al., 2004)

#### Loading and boundary conditions

A uniform pressure load was applied to the inner surface of the corneoscleral shell to simulate the effects of increasing IOP from 0 to 50 mmThe node at the apex of the cornea was constrained in all directions to prevent displacement or rotation, as done previously. (Sigal, 2009, 2011; Sigal and Grimm, 2012) The nodes on the plane of symmetry (**Figure 2E**) were constrained to deform so that they remained on the symmetry plane.*Solutions*. The model was solved using the FEBio software package (Musculoskeletal Research Laboratories, University of Utah, Salt Lake City, UT, USA) with eight-node hexahedral elements. Convergence tests were performed and adequate accuracy (relative strain differences under 1%) was achieved with an average element length of 100 µm. We computed the maximum principal strain as a measure of tissue stretch in response to changes in IOP.

### 2.4 Quantification of collagen fiber recruitment

We tracked the percentage of recruited fibers in each region as IOP increased from 0 to 50 mmHg. We used the model-predicted tissue stretch to determine whether a wavy fiber was recruited. Specifically, if the magnitude of tissue stretch is larger than the tortuosity of a fiber, the fiber is considered recruited; otherwise, the fiber is considered reserved. For each region, we used the number of tortuosity measurements that were smaller than the stretch as an indication of the number of recruited fibers. We then divided this number by the total number of tortuosity measurements in this region, as an indication of the total number of fibers, to calculate the percentage of recruited fibers. The recruitment curve was fitted to the log-normal distribution function using least square method.

## 3. Results

The model-predicted maximum principal strain over the whole globe at an IOP of 50 mmHg is shown in **Figure 3**. The strain distribution was nonuniform throughout the corneoscleral shell. In general, the pattern of strain variations across the shell was similar to that of tortuosity variations shown in **Figure 2C**. For example, regions with large crimp tortuosities experience large strains (i.e., limbus, anterior equator, and equator), and regions with small tortuosities (i.e., posterior equator and posterior sclera) experience small strains.

**Figure 3.**
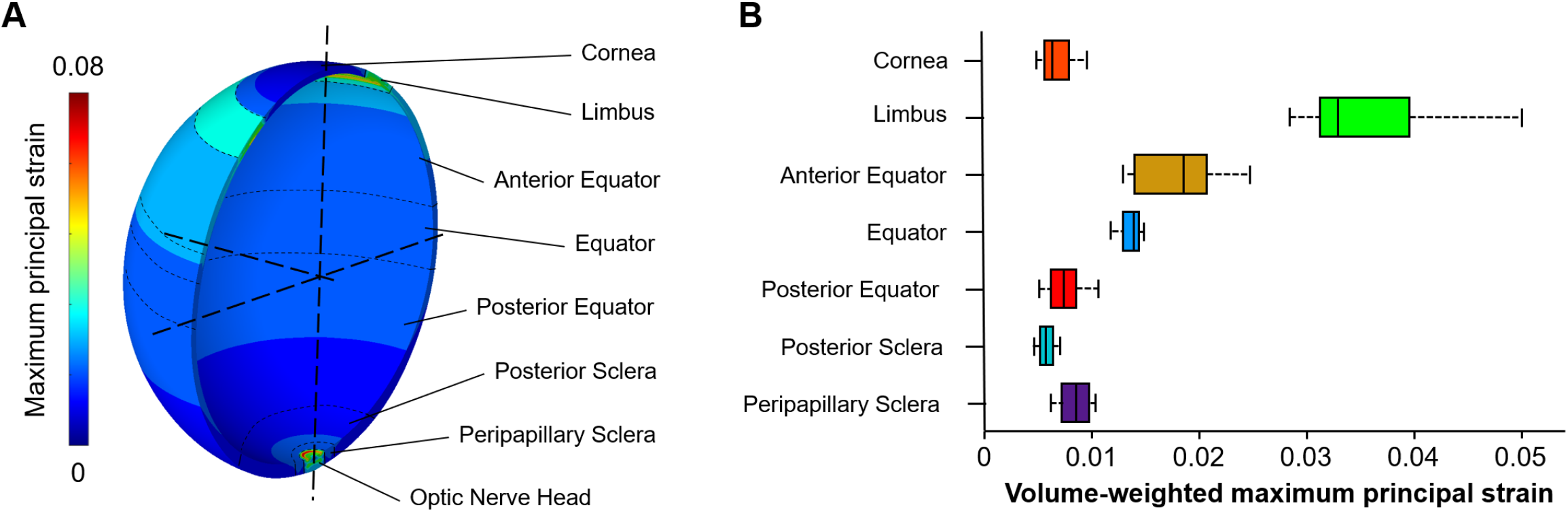
The maximum principal strain over the whole globe at an IOP of 50 mmHg. **(A)** Contour plot of the maximum principal strain. The strain was not uniform across the globe. The highest strain was in the optic nerve head region as this region was more compliant than the other regions. In the corneoscleral shell, the limbus region experienced higher strains than the other regions. **(B)** Box plot of the maximum principal strain in the seven regions across the corneoscleral shell. Note that the strains were weighted by the volume of elements. Overall, the pattern of strain variations across the shell was similar to that of tortuosity variations shown in Figure 2C, but there are also some interesting differences. For example, the equator region had a wide distribution of collagen crimp tortuosity, but the strain distribution was narrow.

Figure 4. shows the contour plots of collagen fiber recruitment over the corneoscleral shell at various IOP levels, with colors corresponding to the percentage of recruited fibers. As IOP increases, collagen fibers of the corneoscleral shell were not recruited simultaneously, suggesting a region-dependent rate of tissue stiffening with IOP. At either normal or elevated IOP, collagen fibers of the corneoscleral shell were not fully recruited. A detailed quantitative analysis of the IOP-induced collagen fiber recruitment over the corneoscleral shell is shown in **Figure 5**.

## 4. Discussion

Our goal was to obtain regionally-resolved estimates of the fraction of corneoscleral collagen fibers recruited, hence contributing to bearing IOP-related loads. The results show that, at low IOPs, collagen fibers in the posterior equator were recruited the fastest, such that at a physiologic IOP of 15 mmHg, over 90% of fibers were recruited, compared with only a third in the cornea and the peripapillary sclera. This, in turn, means that at a physiologic IOP the posterior equator had a fiber reserve of only 10%, whereas the cornea and peripapillary sclera had two thirds. At an elevated IOP of 50 mmHg, collagen fibers in the limbus and the anterior/posterior equator were almost fully recruited, compared with 90% in the cornea and the posterior sclera, and 70% in the peripapillary sclera and the equator. Two main findings can be drawn from these results. First, collagen fibers of the corneoscleral shell were not recruited simultaneously with increasing IOP. Second, collagen fibers of the corneoscleral shell were not fully loaded or recruited at either normal or elevated IOP. Below we discuss each finding in detail.

**Figure 4.**
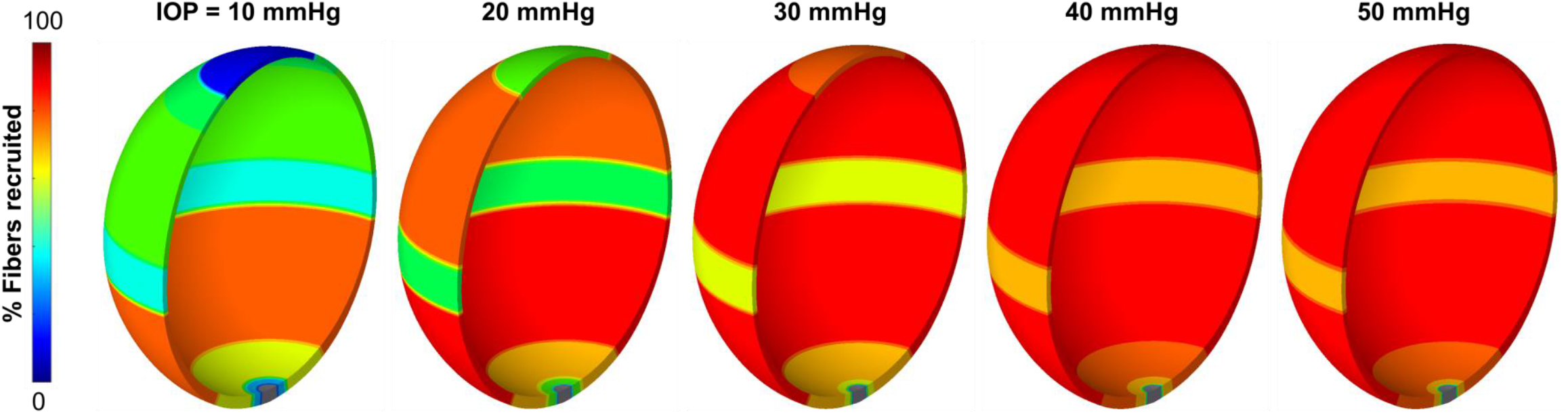
Contour plots of collagen fiber recruitment over the corneoscleral shell at various IOP levels. The shell was colored according to the percentage of recruited fibers, with blue indicating no recruitment and red indicating fully recruitment. Collagen fibers of the corneoscleral shell were not recruited simultaneously, suggesting a region-dependent rate of tissue stiffening with IOP. At either normal or elevated IOP, collagen fibers of the corneoscleral shell were not fully recruited. Note: the scleral canal region is shown in grey since we did not calculate a fiber recruitment for this region.

**Figure 5.**
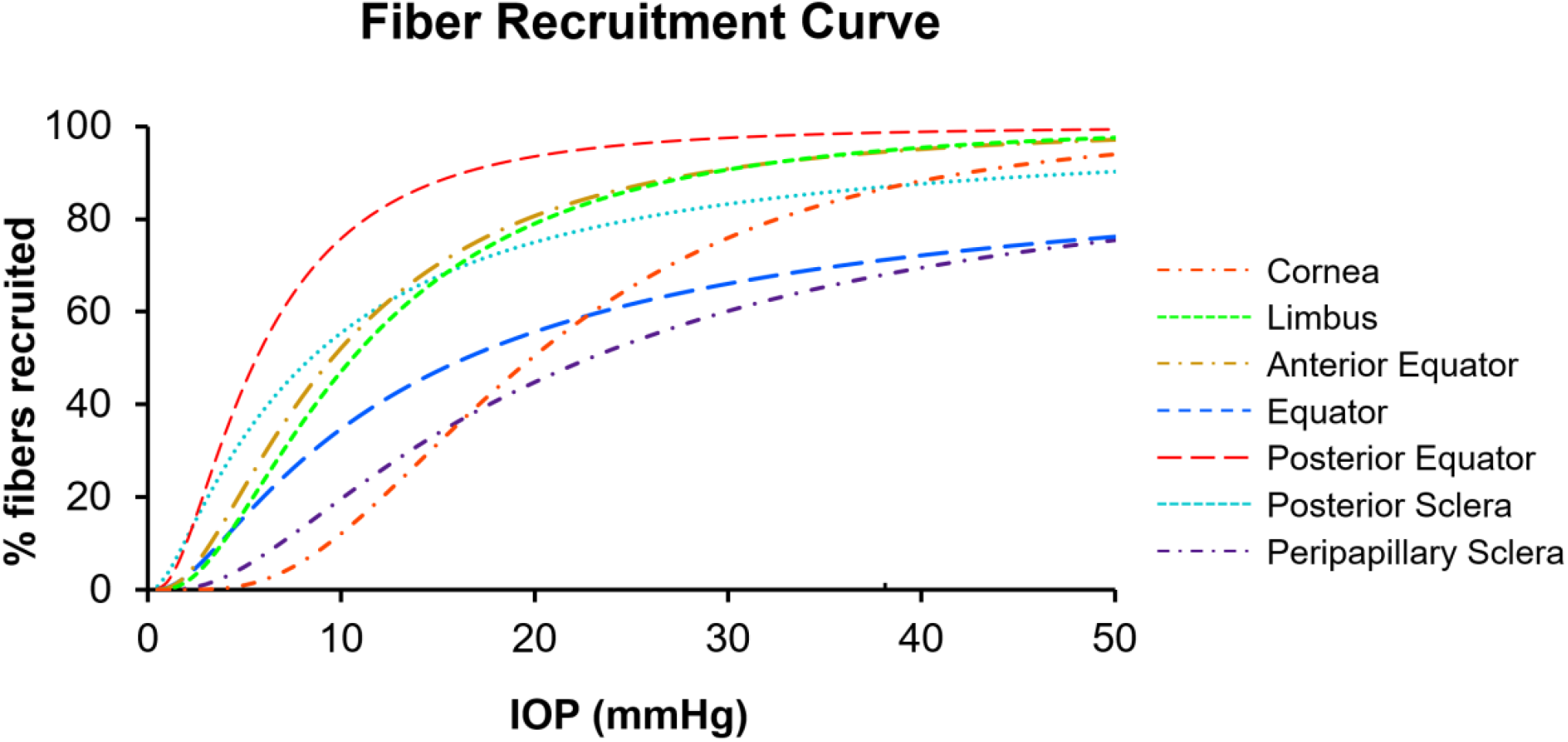
The IOP-induced collagen fiber recruitment over the corneoscleral shell. The curves were colored by regions across the corneoscleral shell. All regions exhibited sigmoid recruitment curves, but the recruitment rates varied substantially. At low IOPs, collagen fibers in the posterior equator were recruited the fastest, such that at a physiologic IOP of 15 mmHg, over 90% of fibers were recruited, compared with only a third in the cornea and the peripapillary sclera. At an elevated IOP of 50 mmHg, collagen fibers in the limbus and the anterior/posterior equator were almost fully recruited, compared with 90% in the cornea and the posterior sclera, and 70% in the peripapillary sclera and the equator.

Our results show that collagen fibers of the corneoscleral shell were not recruited simultaneously with increasing IOP. This suggests that the rate of tissue stiffening with IOP differs in different regions of the corneoscleral shell. We found that collagen fibers in the posterior equator of sclera were recruited the fastest. This could be attributed to the small collagen crimp tortuosity in this region, in which fibers only require a small amount of stretch to straighten. A biomechanical advantage of fast recruitment is to resist large deformations at moderate IOPs, which could help maintain the shape of globe. On the other hand, collagen fibers in cornea were recruited the slowest, suggesting that cornea stiffens at higher IOPs. In this sense, cornea likely behaves as a “mechanical absorber” to protect tissues in other regions from large acute volume or IOP increases. Note that uncrimping or recruitment only relates to the rate of tissue stiffening with IOP, which is different from tissue stiffness. Therefore, our results should not be interpreted to mean that the posterior equator of sclera is stiffer than cornea.

It is important to note that, in addition to collagen crimp, other factors may affect the percentage of collagen recruitment, including, but not limited to, fiber type, composition, alignment, and slip conditions, fiber-to-fiber interactions, proteoglycan and elastin content and distribution, the amount and type of cross-linking between fibers, and the presence of blood vessels within fibers. (Birch et al., 2013; Brazile et al., 2020; Ethier et al., 2004; Fratzl, 2008; Holzapfel, 2001; Wang et al., 2020) Further research is needed to quantify additional microstructural characteristics to be able to fully characterize the IOP-induced collagen fiber recruitment over the corneoscleral shell.

Our results also show that collagen fibers of the corneoscleral shell were not fully loaded or recruited at either normal or elevated IOP. Note that collagen fibers in other connective tissues, such as artery, (Hill et al., 2012; Roy et al., 2010; Weisbecker et al., 2015) heart valve, (Mattson et al., 2017) and bladder, (Cheng et al., 2018) are also not fully recruited under physiological loading. However, most models developed to study connective tissue mechanics only consider the fiber density, orientation, and degree of anisotropy or fiber concentration, assuming all fibers are recruited or loaded. (Coudrillier et al., 2013; Hua et al., 2020; Pandolfi and Holzapfel, 2008; Thomas et al., 2019; Voorhees et al., 2017c; Zhang et al., 2015; Zhou et al., 2019a) This may overestimate the stiffness of connective tissues, resulting in a relatively low physiological accuracy of the estimates obtained from models.

We understand that an IOP of 50 mmHg is high. In our study, we aimed to investigate the biomechanics of the eye under extreme conditions and provide insight into how the eye responds to different mechanical stresses. Moreover, it is not uncommon for studies in ocular biomechanics, particularly those related to ONH biomechanics, to consider high or extremely high pressures, such as those that reach 45 or 50 mmHg. (Amini et al., 2011; Brazile et al., 2020; Gsellman and Amini, 2016; Ho et al., 2014; Sigal et al., 2004; Sigal et al., 2009a; Wang et al., 2016b) By extending our work to such pressures, we intended to relate to those studies and provide further insight into the biomechanics of the eye under various conditions. Interestingly, even at the extremely high IOP of 50 mmHg, not all fibers are recruited. This suggests that some fibers may be optimized to bear forces that are not due to IOP, such as from intracranial pressure, (Feola et al., 2016; Feola et al., 2018; Hua et al., 2017; Hua et al., 2018) gaze, (Sibony et al., 2018; Wang et al., 2016a; Wang et al., 2016b) muscle tension, (Jafari et al., 2021) or impact. This conclusion would not be possible if we had only considered normal or mildly elevated IOPs.

It should be noted that the peripapillary sclera recruitment curve as a function of IOP derived in this study is generally lower than that observed in our previous study. (Jan and Sigal, 2018) For example, at a normal IOP, this study observed approximately 1/3 recruitment, compared to 75% in the previous study. Interestingly, however, at a high IOP of 50 mmHg, both studies showed similar results with about 70% recruitment. It is important to keep in mind that the differences and similarities between the two studies should be interpreted with caution due to several factors that could have impacted the results. First, in the two studies different species were investigated: in the previous one sheep, and in this one humans. Although the species are similar, it cannot be assumed that the results will be identical. Second, the direction of measurement is different, with the previous study using coronal sections and this study using longitudinal sections. This difference could potentially impact the visibility of crimp and recruitment as the measurements represent fibers aligned with the plane of sectioning. Third, in the previous study we directly measured crimp from eyes fixed at various IOPs, whereas in this study we used crimp from eyes at no pressure to estimate the recruitment resulting from increases in IOP. As noted elsewhere, fiber recruitment is not the only factor affecting tissue stiffness. Since we have not incorporated all these factors, predictions made in this study may be off. Fourth, in the microstructural model used in this study, we assumed a monotonic increase in the fraction of recruitment with IOP. In contrast, in the previous study, we observed a maximum recruitment fraction of 82% at approximately 25 mmHg of IOP, followed by a decrease to 75% at 50 mmHg.

The relationship between collagen microstructure and the overall tissue mechanical properties is complex. (Amini et al., 2014; Thomas et al., 2019) We developed a fiber-based microstructural constitutive model that considered crimp tortuosity of collagen fibers. Collagen fibers were assumed as linear elastic, and the nonlinear stiffening behavior was derived from a load-dependent recruitment of collagen fibers based on a distribution of crimp tortuosities. A similar approach has been adopted by others. (Cacho et al., 2007; Liu et al., 2014; Raz and Lanir, 2009) Inversely, Grytz et al. predicted collagen crimp morphology in the corneoscleral shell using the stress-strain curves gathered from mechanical tests. (Grytz and Meschke, 2009, 2010) They assumed that collagen fibers in the corneoscleral shell had uniform helical spring morphology and were never straightened under stretch. In their constitutive model, the nonlinearity of the stress-strain curve arose from fiber uncrimping, rather than recruitment. These are related but not the same. We found direct experimental evidence that the peripapillary sclera gradually recruited its collagen fibers with increasing IOP-induced stretch. (Jan et al., 2017a; Jan and Sigal, 2018; Lee et al., 2022)

We used tortuosity, i.e., the path length divided by the end-to-end length (≥ 1), as a measure of crimp morphology in this study. When the tortuosity of a fiber drops to one under stretch, the fiber is considered recruited. It is important to note that tortuosity is only one measure of crimp morphology. Other measures, such as period (the length of one wave), waviness (the standard deviation of orientations), and amplitude (half the peak to trough distance), can also be used to quantify crimp morphology, and could vary in different ways than tortuosity. (Gogola et al., 2018a; Jan et al., 2018; Jan et al., 2017a; Jan and Sigal, 2018) Future work should incorporate additional crimp measures in the constitutive model to fully characterize the relationship between collagen crimp morphology and the nonlinear mechanical behavior of tissues.

The crimp tortuosity incorporated in our constitutive model was quantified using PLM. (Gogola et al., 2018a) Others have visualized the crimp in the eye with other imaging modalities, including transmitted electron microscopy, (Liu et al., 2014) brightfield microscopy, (Ostrin and Wildsoet, 2016) nonlinear microscopy, (Quantock et al., 2015) and MRI. (Ho et al., 2014) The advantages of PLM have been detailed in our previous work. (Gogola et al., 2018a; Jan et al., 2018; Jan et al., 2017a; Jan and Sigal, 2018; Jan et al., 2015) Briefly, PLM has the appropriate sensitivity, resolution, and field of view to quantify crimp over the whole globe. In addition, sample preparation for the crimp analysis is relatively simple as tissue can be visualized without dehydration, labels, or stains, reducing tissue processing and the likelihood of introducing artifacts, such as deformation.

The three-dimensional axisymmetric model presented in this study is a simplification of the corneoscleral shell. It follows the general idea of generic parametric models we have published previously. (Hua et al., 2017; Hua et al., 2018; Sigal, 2009; Sigal et al., 2005a; Sigal et al., 2004; Sigal and Grimm, 2012; Sigal et al., 2011a; Sigal et al., 2011b; Voorhees et al., 2016) Although the simplifications may not capture every aspect of the complex mechanical behavior of the corneoscleral shell, they are well-suited to test the fundamental behavior of the parameters of interest, without the overlying complexity of inter-eye variability that the specimen-specific models imply. (Sigal et al., 2005b; Sigal et al., 2009a, b; Sigal et al., 2010a; Sigal et al., 2010b; Voorhees et al., 2020; Voorhees et al., 2017b; Voorhees et al., 2017c) In addition, these simplifications help provide fundamental new insight into how collagen fibers of the corneoscleral shell were recruited with increasing IOP, which would otherwise be challenging to obtain through experiment. A thorough discussion of the limitations of this modeling approach can be found in our previous studies, (Hua et al., 2017; Hua et al., 2018; Sigal, 2009; Sigal et al., 2005a; Sigal et al., 2004; Sigal and Grimm, 2012; Sigal et al., 2011a; Sigal et al., 2011b; Voorhees et al., 2016) and was recently discussed in detail. (Roberts et al., 2018) Below we summarize the limitations more relevant to this study.

First, the crimp tortuosity incorporated in our constitutive model was measured in tissues that had been histologically processed. (Gogola et al., 2018a) Some artifacts, such as tissue distortion or shrinkage, could result from fixation or sectioning. However, because all the globes were treated the same way, the regional differences in crimp tortuosity as we measured must still be valid. Further studies could measure crimp in tissues fixed with 10% formalin, as it has shown to have minimal effects on the size or shape of ocular tissues at large and small scales, (Jan et al., 2015; Tran et al., 2017) or study crimp without sectioning. (Ho et al., 2014)

Second, we measured the crimp tortuosity based on the in-plane collagen fiber orientations derived from 2D histological sections of the corneoscleral shell. (Gogola et al., 2018a) Although we were careful to measure collagen fibers and bundles in the section plane, it is likely that they had small out-of-plane angles. In this case, our 2D measures of crimp tortuosity would be underestimated relative to the three-dimensional variations. Future studies should include examination of any potential differences with three-dimensional measurements. This examination could be done, for example, by using sections made with different orientations, or using extensions of PLM that allow three-dimensional measurement of fiber orientation. (Yang et al., 2018)

Third, our constitutive model did not consider the elastic responses of cells, proteoglycans, elastin, glycosaminoglycans, and other extracellular tissue constituents. (Hatami-Marbini and Pachenari, 2020; Midgett et al., 2020; Murienne et al., 2016) As a result, the contribution of these constituents to the tissue mechanical properties or the interaction of these constituents with the collagen fiber network was ignored in our constitutive model. We remind readers that our intention was not to accurately quantify the IOP-induced deformation of the globe, but to understand how collagen fibers of the corneoscleral shell bear loads as IOP increases. In this sense, it is acceptable for our constitutive model to focus on the elastic response of collagen fibers.

Fourth, collagen fibers of the corneoscleral shell are interwoven. (Gogola et al., 2018b; Jan et al., 2017b; Komai and Ushiki, 1991; Lee et al., 2022; Meek, 2009; Morishige et al., 2007; Nguyen, 2016) We and others have demonstrated that collagen interweaving plays an important role in determining the structural stiffness of ocular connective tissues. (Pandolfi et al., 2019; Petsche and Pinsky, 2013; Wang et al., 2020; Winkler et al., 2011) However, such interweaving was not considered in our constitutive model. Future constitutive models might benefit from incorporating measures of collagen interweaving from experiments to better characterize the IOP-induced collagen fiber recruitment over the corneoscleral shell.

Fifth, our finite element model did not incorporate depth-dependent variations in fiber distributions. Using wide-angle x-ray scattering, Pijanka et al. found that fiber concentration factors varied with depth. (Pijanka et al., 2012) Another study by Danford et al., using small angle light scattering, confirmed depth dependence of the fiber concentration factor in both normal and glaucoma human sclera. (Danford et al., 2013) More advanced finite element models will be required to understand the implication of depth-dependent fiber concentration factors on the IOP-induced globe deformation and collagen recruitment.

Sixth, our model did not incorporate prestresses.(Grytz and Downs, 2013) It is unclear how exactly prestresses would change the conclusions, especially since the magnitudes of the prestresses likely also vary over the globe and are anisotropic. Although it has been shown that residual strains in soft tissues can be measured experimentally, (Amini et al., 2012; Wang et al., 2015) to the best of our knowledge the techniques have yet to be applied over the whole globe.

Seventh, the experimental data on collagen fiber tortuosity on which we based this study only accounted for variations along the axial length of the globe, aggregating measurements over equatorial lines. Hence the models and analysis in this work were axisymmetric, which may be useful as a first approximation, (Hua et al., 2018) but it also means that this work cannot account for more complex conditions. (Chung et al., 2016) More detailed experimental information on the anisotropy of the sclera, both in terms of its mechanical behavior and collagen components, will be necessary for more comprehensive models. (Hua et al., 2022; Jan et al., 2022)

Eighth, we modeled the ONH tissues as linearly elastic. We did this for simplicity to avoid more complex boundary conditions that could affect the interpretation of our results, which were primarily focused on the sclera.

Lastly, experimental measurement of whole-globe collagen fiber recruitment with IOP is not possible yet with current imaging techniques, making it difficult to validate our model predictions. With the development of more advanced imaging techniques, such as polarization-sensitive optical coherence tomography, (Baumann et al., 2014; Tang et al., 2021; Willemse et al., 2020) we expect relevant experimental data to be available in the near future.

In conclusion, by developing a fiber-based microstructural constitutive model that could account for the collagen fiber crimp via their tortuosity, we obtained regionally-resolved estimates of the fraction of corneoscleral collagen fibers recruited and in reserve. An accurate measure of the fraction of the fibers bearing load can help improve our understanding and ability to predict and control tissue remodeling and/or treatments based on fiber cross-linking.

## Acknowledgements

Supported in part by National Institutes of Health grants R01-EY023966, R01-EY028662, P30-EY008098, and T32-EY017271; Eye and Ear Foundation (Pittsburgh, PA); Research to Prevent Blindness (unrestricted grant to UPMC Ophthalmology and Stein Innovation Award to Sigal IA); BrightFocus Foundation; National Science Foundation (CAREER 2049088).

