## Supplementary Material for "Who bears the load? IOP-induced collagen fiber recruitment over the corneoscleral shell"

**Short title:** Collagen recruitment over the whole globe

**† Correspondence:**
Ian A. Sigal, Ph.D.
Laboratory of Ocular Biomechanics
Department of Ophthalmology, University of Pittsburgh School of Medicine
203 Lothrop Street, Eye and Ear Institute, Rm. 930, Pittsburgh, PA 15213

www.OcularBiomechanics.com

**Keywords**: collagen, crimp, tortuosity, cornea, sclera, recruitment, biomechanics

**Disclosures:** None.

**Funding:** Supported in part by National Institutes of Health grants R01-EY023966, R01-EY028662, P30-EY008098, and T32-EY017271; Eye and Ear Foundation (Pittsburgh, PA); Research to Prevent Blindness (unrestricted grant to UPMC Ophthalmology and Stein Innovation Award to Sigal IA); BrightFocus Foundation; National Science Foundation (CAREER 2049088).

The strain energy function of the sclera was defined as

$$W\,(\boldsymbol{C}) = W_{\text{matrix}} + \int_{\theta_p-\frac{\pi}{2}}^{\theta_p+\frac{\pi}{2}} P(\theta) W_{fiber}\big(I_4(\theta)\big) d\theta$$

The ground substance matrix of the sclera was modeled as a Neo-Hookean material such as

$$W_{\text{matrix}}(I_1) = c_1(I_1 - 3)$$

The strain energy associated with the collagen fiber family was defined as

$$W_{fiber} = \begin{cases} 0 & (\text{when } \lambda < \lambda_m) \\ \xi\left(\lambda_m \ln\left(\frac{\lambda_m}{\lambda}\right) + (\lambda - \lambda_m)\right) & (\text{when } \lambda \geq \lambda_m) \end{cases}$$

For the convexity requirement, the strain energy function should satisfy the equation below:

$$\frac{\partial^2 W(\boldsymbol{C})}{\partial \boldsymbol{C}^2} \geq 0$$

When $\lambda < \lambda_m$, $W_{fiber} = 0$. So $W\,(\boldsymbol{C}) = W_{\text{matrix}}$, $\frac{\partial^2 W(\boldsymbol{C})}{\partial \boldsymbol{C}^2} = 0$ fulfilled the convexity requirement.

When $\lambda \geq \lambda_m$, without considering von-Mises distribution (set $\theta_p = 0,\ k \longrightarrow \infty$),

$$W(\boldsymbol{C}) = W_{\text{matrix}}(I_1) + W_{fiber}(I_4)$$

where $W_{\text{matrix}}(I_1) = c_1(I_1 - 3)$,$W_{fiber}(I_4) = \xi\left(\lambda_m \ln\left(\frac{\lambda_m}{\lambda}\right) + (\lambda - \lambda_m)\right)$.

$$\frac{\partial^2 W(\boldsymbol{C})}{\partial \boldsymbol{C}^2} = \frac{\partial}{\partial I_1}\left(\frac{\partial W_{\text{matrix}}(I_1)}{\partial I_1}\frac{\partial I_1}{\partial \boldsymbol{C}}\right)\frac{\partial I_1}{\partial \boldsymbol{C}} + \frac{\partial}{\partial I_4}\left(\frac{\partial W_{fiber}(I_4)}{\partial I_4}\frac{\partial I_4}{\partial \boldsymbol{C}}\right)\frac{\partial I_4}{\partial \boldsymbol{C}}$$

$$= 0 + \frac{\partial}{\partial I_4}\left(\frac{\partial W_{fiber}(I_4)}{\partial I_4}\frac{\partial I_4}{\partial \boldsymbol{C}}\right)\frac{\partial I_4}{\partial \boldsymbol{C}} = \frac{\partial}{\partial I_4}\left(\frac{\xi(\lambda - \lambda_m)}{2\lambda^2}\boldsymbol{a}\otimes\boldsymbol{a}\right)\frac{\partial I_4}{\partial \boldsymbol{C}}$$

$$= \frac{\partial}{\partial I_4}\left(\frac{\xi\left(\lambda_m - {I_4}^{-1/2}\right)}{2{I_4}^2}\boldsymbol{a}\otimes\boldsymbol{a}\right)\frac{\partial I_4}{\partial \boldsymbol{C}} = \frac{\xi(2\lambda_m - \lambda)}{4\lambda^4}\boldsymbol{a}\otimes\boldsymbol{a}\otimes\boldsymbol{a}\otimes\boldsymbol{a}$$

When $2\lambda_m - \lambda > 0$, i.e., $\lambda < 2\lambda_m$, the convexity was fulfilled.
